# Neutrophils matter: New clinical insights on their role in the progression of metastatic breast cancer

**DOI:** 10.1101/2025.04.11.648384

**Authors:** Bruna F. Correia, Daniela Grosa, Rute Salvador, Inês Brites, Telma Martins, Marina Vitorino, Carolina Xavier Sousa, Sofia Cristóvão-Ferreira, Sofia Braga, António Jacinto, Maria Guadalupe Cabral

**Author notes:** These authors contributed equally to this work (Bruna F. Correia, Daniela Grosa).

## Abstract

**Background:** Metastatic breast cancer (BC) remains a significant clinical challenge, needing innovative strategies to improve disease management and extend patient survival. Increased neutrophil levels have been observed in both peripheral blood and tumor tissues of patients with different types of cancer, often being associated with poor clinical outcomes. These findings suggest a crucial role for neutrophils in tumor progression, raising interest in neutrophil-based therapies. However, the functional and phenotypic heterogeneity of neutrophils complicates their therapeutic targeting. This study aims to investigate the clinical impact of immunosuppressive, protumor low-density neutrophils (LDN) in metastatic BC, comparing them with normal high-density neutrophils (HDN) to better understand their role in disease progression.

**Methods:** LDN and HDN subpopulations were isolated from the blood of 151 BC patients (72 metastatic, 79 non-metastatic) using density gradient centrifugation. Their frequency, phenotype, and function were analyzed by flow cytometry and *in vitro* experiments. Correlations between LDN levels and clinical data from metastatic BC patients were evaluated, alongside individual longitudinal assessments.

**Results:** LDN accumulated significantly in the blood of BC patients, particularly in those with metastatic disease. Elevated LDN levels in these patients were associated with faster disease progression and reduced life expectancy, regardless of metastatic site. Longitudinal analysis revealed that higher LDN percentages often correlated with adverse clinical events, whereas lower levels of LDN were linked to stable disease. Functionally, LDN exhibited protumor properties, including elevated expressions of PD-L1 and MMP-9, contributing to immunosuppression and metastasis. Unlike HDN, which demonstrated cytotoxicity against tumor cells, LDN failed to reduce BC cell line viability in 3D co-cultures. Notably, BC cell lines exposed to LDN-conditioned medium showed increased invasive capacity and proliferation, while T cells cultured in the same medium displayed impaired activation, likely due to the effect of arginase.

**Conclusion:** Our results highlighted LDN and their secreted factors as major drivers of BC progression and increased aggressiveness. These findings suggest that incorporating LDN assessment into clinical practice could aid in identifying high-risk patients and enable more personalized treatment approaches. Furthermore, our data strengthen the relevance of targeting specific neutrophil subsets or their functions to improve metastatic BC management and patient outcomes.

## Background

Breast cancer (BC) is the most prevalent malignancy and the leading cause of cancer-related mortality among women worldwide [1], with metastatic BC (mBC) responsible for the vast majority of these deaths [2]. Despite advances in cancer treatment, managing metastatic BC (mBC) remains a major challenge, with systemic chemotherapy as the primary therapeutic option to control disease progression and extend patient survival [3]. However, the development of chemotherapy resistance significantly limits treatment efficacy and reduces survival rates in affected patients [4]. Therefore, elucidating the mechanisms underlying this resistance is essential for optimizing therapeutic strategies and improving outcomes in mBC patients.

The tumor microenvironment (TME) exerts complex and context-dependent effects on cancer progression, largely driven by tumor-infiltrated immune cells that promote immunotolerance and support tumor growth [5]. Among these immune components, neutrophils have recently emerged as key contributors to tumor progression and mediators of therapy resistance [6,7,8,9]. For instance, analysis of a human BC dataset revealed a higher expression of neutrophil-related signature genes in tumors from therapy-resistant patients compared with therapy-sensitive ones [10]. This finding highlights the potential role of neutrophils in modulating therapeutic outcomes.

Once considered passive bystanders in cancer biology, neutrophils are now recognized as a heterogeneous population of cells with remarkable phenotypic and functional diversity, capable of shaping tumor behavior [6,11]. Within the TME, at least two distinct neutrophil populations have been identified, N1 and N2, exhibiting antitumor and protumor properties, respectively [12,13,14].

This renewed understanding of neutrophil biology has opened new possibilities for expanding cancer therapies. However, significant challenges remain, particularly in distinguishing between harmful and beneficial neutrophil populations, making it difficult to selectively target or modulate these cells without unintended consequences.

Systemically, similar to the N1 and N2 neutrophil subsets identified within the TME, two distinct populations of circulating neutrophils have been identified, each playing different roles in cancer progression. Their functional and phenotypic diversity in the bloodstream allows their isolation via density gradient centrifugation, separating them into two fractions: high-density neutrophils (HDN) and low-density neutrophils (LDN) [15,16,17]. HDN, considered the normal neutrophils, exhibit a pro-inflammatory phenotype and cytotoxic activity, supporting antitumor immune responses and counteracting cancer progression [16,17,18,19]. In contrast, LDN display immunosuppressive and protumor properties, facilitating cancer growth and metastasis [16,17,20,21]. LDN represent a heterogeneous population comprising both mature and immature neutrophils, predominantly emerging under inflammatory or pathological conditions and rarely found in the blood of healthy individuals [16,17,22,23]. Their presence in disease states has sparked significant clinical interest.

Mechanistically, LDN are implicated in tumor progression through immunosuppressive functions, including the secretion of ROS, NO, and arginase, PD-L1 expression, as well as the stimulation of regulatory T cells, all of which inhibit effector T lymphocyte activity [16,24,25]. Additionally, LDN promote metastasis formation through the release of metalloproteinases and neutrophil extracellular traps (NETs), facilitating extracellular matrix degradation and the migration of circulating tumor cells to secondary niches [20,26]. Despite growing interest, research on neutrophil functions in cancer remains relatively new, with most existing studies on their mechanisms relying on murine models, which limits their direct relevance to human disease. Thus, while some clinical studies have been conducted, there is a pressing need for further investigations using human samples for a more comprehensive understanding of the neutrophil roles in cancer biology. Longitudinal studies are particularly relevant to systematically assess whether LDN can serve as biomarkers or predictors of high-risk patients, disease aggressiveness, and treatment resistance.

To address these gaps and further validate the functional significance and clinical impact of LDN our team has been studying neutrophils derived from BC patients. Previously, we demonstrated that systemic LDN levels are significantly elevated in the blood of BC patients compared to healthy individuals and are strongly associated with poor responses to neoadjuvant chemotherapy and metastatic disease [16]. Building on these findings, we aimed to expand our analysis by studying a larger cohort of BC patients, with a particular focus on advanced disease. Additionally, we performed individual follow-up analyses to explore LDN relevance in disease progression.

Our results confirm that LDN accumulate significantly in the blood of metastatic BC patients and that higher LDN levels in these patients are linked to accelerated disease progression and reduced survival probability, regardless of the metastatic site. To our knowledge, this is the first BC longitudinal study to demonstrate that increased LDN percentages frequently correlate with adverse clinical outcomes, while lower LDN levels are associated with stable disease. Furthermore, functional assays revealed that not only LDN but also their conditioned media alone exhibit strong immunosuppressive and protumor properties, further highlighting their critical role in BC progression.

These findings underscore the potential of LDN as both prognostic biomarkers and novel therapeutic targets, offering promising avenues to improve the management and outcomes of BC patients.

## Material and Methods

### Patients’ samples

Blood samples were collected from 151 breast cancer (BC) patients, including 79 non-metastatic and 72 metastatic cases. Whenever possible, follow-up samples were also collected, approximately every 3 months, for the 48 metastatic BC patients included in the longitudinal study (see flowchart in Supplementary Figure 1). Additionally, whole blood from 8 healthy donors was collected for comparison. All blood samples were collected in Vacutainer EDTA tubes (BD Biosciences) and processed within 24 hours post-collection.

Patients’ clinical and demographic characteristics are detailed in Table 1. Samples were provided by six hospitals: Unidade Local de Saúde Amadora Sintra – Hospital Professor Doutor Fernando Fonseca (HFF), Unidade Local de Saúde do Arco Ribeirinho – Hospital de Nossa Senhora do Rosário (ULSAR), Hospital de Vila Franca de Xira (HVFX), Hospital Santa Maria (HSM), Hospital CUF Descobertas (HCD) and Instituto Português de Oncologia de Lisboa Francisco Gentil (IPO).

**Table 1.** Characteristics of the non-metastatic and metastatic breast cancer patients enrolled in the study. Summary of age, body mass index, and clinical characteristics, including primary tumor breast cancer subtype, size and Ki67 percentage, and metastasis location, of the patients enrolled in this study. Clinical data were not available for all participants, with primary tumor information partially missing for patients who were not under the current hospital follow-up at the time of initial diagnosis.

|  | Patients |  |
| --- | --- | --- |
|  | Non-metastatic | Metastatic |
| Number of subjects | 79 | 72 |
| Age | Median: 54.12<br>(Range: 31 - 80) | Median: 57.28<br>(range: 30 - 84) |
| Body Mass Index (BMI) | Median: 27.78<br>(range: 17.22 - 46.68) | Median: 25.88<br>(range: 17.63 – 38.75) |
| Primary breast cancer subtype: |  |  |
| Luminal A | 5 | 16 |
| Luminal B | 10 | 38 |
| HER2+ | - | 8 |
| TNBC | 10 | 7 |
| Primary tumor dimension (mm) | Median: 38.97<br>(range: 6 - 110) | Median: 49.70<br>(range: 9 - 280) |
| % of Ki67 | Median: 49.22%<br>(range: 5 - 98%) | Median: 31.20%<br>(range: 2 - 100%) |
| Metastasis location: |  |  |
| Bone | NA | 46 |
| Brain | NA | 10 |
| Liver | NA | 32 |
| Lung | NA | 26 |
| Others | NA | 42 |

All participants had breast tumors or metastasis derived from a primary breast tumor, were over 18 years of age, and were able to understand the study in which they were involved. Samples were collected as part of the routine clinical care and did not interfere with patients’ treatment or diagnosis.

### Sample processing

Low-density neutrophils (LDN) and high-density neutrophils (HDN) were isolated from whole blood using a Histopaque-based density gradient centrifugation method. Briefly, whole blood was layered on top of a solution of equal volumes of Histopaque-1119 and Histopaque-1077 (Sigma-Aldrich), in a 1:1 ratio (blood:Histopaque solution) and centrifuged at 2000 rpm for 20 minutes, without applying a brake. After centrifugation, circulating LDN, which may emerge in the peripheral blood mononuclear cell (PBMC) layer, and normal/HDN, which are found in the granulocyte fraction, were collected. The frequency of LDN and HDN was then assessed in the PBMC and granulocyte layers, respectively, and further immunophenotyping of both populations was performed by flow cytometry. The LDN frequency was then compared with patients’ clinical data.

Neutrophils were further enriched from each cell fraction by positive selection of CD15+ cells using the human CD15 MicroBeads Kit (Miltenyi Biotec) and LS columns to ensure high purity. The enriched neutrophil populations (LDN and HDN) were subsequently used in functional *in vitro* assays.

### Immunophenotyping by flow cytometry

Antibody staining was performed on whole blood and on LDN and HDN fractions. A cocktail of mouse anti-human monoclonal fluorescent antibodies (mAbs) against selected cell surface molecules was added to the samples and incubated in the dark for 15 minutes at room temperature. For both whole blood and HDN fractions, red blood cell lysis was carried out with RBC lysis buffer (BioLegend), for 20 minutes at 4°C, followed by a wash step with 1X PBS and centrifugation at 300 g for 5 minutes. For intracellular staining, cells were fixed and permeabilized using a fixation/permeabilization solution (Invitrogen) according to the manufacturer’s instructions. Permeabilized cells were incubated with the intracellular mAbs for 30 minutes in the dark at room temperature. In the cases where these antibodies were not conjugated (namely, anti-MPO, and anti-VEGF), a secondary antibody, Alexa Fluor 488 (Life Technologies), was added afterward and incubated for another 30 minutes, followed by a washing step.

Data acquisition was performed using a BD FACS Canto II equipped with FACSDiva Software v8.0.1 (BD Biosciences), and results were analyzed with FlowJo software v10. The mAbs used for the staining of cell surface markers were: anti-CD3-APC (clone UCHT1), anti-CD4-FITC (clone OKT4), anti-CD8-PE (clone HIT8a), anti-CD8-Pacific Blue (clone HIT8a), anti-CD11b-FITC (clone ICRF44), anti-CD14-FITC (clone 63D3), anti-CD15-PE (clone H198), anti-CD25-PE (clone BC96), anti-CD33-APC-Cy7 (clone P67.6), anti-CD66b-APC (clone G10F5), anti-CD69-PercP (clone FN50), anti-CD127-PE-Cy7 (clone A019D5), anti-PD-L1-APC (clone29E.2A3), and anti-TLR4-PE (clone HTA125) all from BioLegend. The mAbs used for the staining of intracellular markers were: anti-MMP-9-FITC (sc-393859), anti-Neutrophil Elastase-AlexaFluor647 (sc-55549), anti-MPO (sc-52707) and anti-VEGF (sc-7269), all from Santa Cruz Biotechnology.

The different immune populations were defined as follows: LDN (in the PBMC fraction), HDN (in the granulocytes fraction), and total neutrophils (in whole blood) as CD15+/CD14-, cytotoxic T lymphocytes as CD3+/CD8+, helper T lymphocytes as CD3+/CD4+ and regulatory T cells as CD4+/CD25high/CD127low; all immune populations are represented as a percentage within the single cells’ gate. Positive populations were established considering the unstained controls.

### Cell lines

*In vitro* experiments were performed using the triple-negative breast cancer (TNBC)-derived cell line MDA-MB-231. The cells were cultured in Dulbecco’s Modified Eagle Medium (DMEM, Gibco) supplemented with 10% fetal bovine serum (FBS, Biowest) and 1% Penicillin/Streptomycin (GE Healthcare). MDA-MB-231 cells were maintained as monolayer cultures in T75 flasks under humidified conditions at 37°C with 5% CO_2_. Once cells reached 80–90% confluency they were passaged by detaching them with TrypLE Express (Gibco) and reseeding them as required for subsequent experiments.

### Cytotoxicity assay in 3D-co-cultures

The MDA-MB-231 BC cell line was cultured in DMEM supplemented with 10% FBS and 1% Penicillin/Streptomycin. Three-dimensional (3D) co-cultures of MDA-MB-231 cells with either patient-derived LDN or HDN were established in agarose-coated plates at a 1:3 ratio (cancer cells:neutrophils), adapting a protocol previously established by our group [27]. After 24 hours of incubation at 37°C, spheroids were harvested, dissociated into single cells by pipetting, and stained with a Fixable Viability Dye (BD Biosciences) and a pan-leukocyte marker, anti-CD45 (clone HI30, BioLegend), to distinguish tumor cells from immune cells. The percentage of viable cancer cells within the CD45 negative population was then evaluated by flow cytometry.

### Invasion assay with neutrophil-conditioned medium

Neutrophils were isolated as previously described, followed by washing, red blood cell lysis, and cell counting. LDN and HDN were enriched from the corresponding fractions using magnetic beads, following the manufacturer’s protocol, and subsequently cultured in RPMI 1640 (Gibco) supplemented with 10% FBS and 1% Penicillin/Streptomycin, for 24 hours at 37°C, 5% CO₂. MDA-MB-231 cells were concurrently incubated in serum-free (SF) medium supplemented with 0.5% bovine serum albumin (BSA) and 1% Penicillin/Streptomycin, under the same conditions.

After 24 hours, MDA-MB-231 cells were resuspended in either neutrophil-conditioned medium or SF medium (as control) and seeded at a density of 0.5 × 10⁵ cells into the upper chambers of Matrigel®-coated transwell inserts (Corning, 8 μm pore size). The inserts were placed in wells containing RPMI with 10% FBS and 1% Penicillin/Streptomycin and cells were allowed to adhere and invade through the Matrigel for 18 hours under humidified conditions at 37°C with 5% CO2.

Following incubation, non-invading cells remaining in the upper chamber were gently removed using a sterile cotton swab. Cells that had invaded through the Matrigel-coated membrane were fixed with 4% paraformaldehyde (PFA) for 15 minutes at room temperature and stained with DAPI (1 µg/mL) for 10 minutes, protected from light. The transwell membranes were carefully excised and mounted onto microscope slides.

Images of the invaded cells were captured using a Zeiss Z2 microscope (20x objective), acquiring 10 random fields per condition. The number of invaded cells was quantified using Fiji software (ImageJ).

### Proliferation assay with neutrophil-conditioned medium

Neutrophils were isolated as previously described, and HDN and LDN were enriched using magnetic beads according to the kits’ protocol. These cells were then cultured in RPMI 1640 supplemented with 10% FBS and 1% Penicillin/Streptomycin, with plating conditions adjusted to cell number, for 24 hours at 37°C in a 5% CO₂ incubator. In parallel, MDA-MB-231 cells were seeded at a density of 5 × 10^4^ cells per well in a 24-well plate in RPMI supplemented with 10% FBS and 1% Penicillin/Streptomycin. After 24 hours, the culture medium was replaced with HDN or LDN-conditioned medium, and cells were incubated for an additional 24 hours at 37°C in a 5% CO₂ incubator. Following this incubation, the MDA-MB-231 cells were detached and stained with the proliferation marker Ki67 (clone Ki-67, BioLegend). The percentage of Ki67+ proliferating cancer cells was then evaluated by flow cytometry.

### T-cell activation assay with neutrophil-conditioned medium

HDN, LDN, and PBMC were isolated from whole blood using density gradient centrifugation, followed by magnetic beads-based sorting according to the manufacturer’s protocol. The cells were cultured in RPMI 1640 supplemented with 10% FBS and 1% Penicillin/Streptomycin, with plating conditions adjusted based on cell numbers. Part of the LDN and HDN cultures was further incubated with the arginase inhibitor nor-NOHA (1 mM, Sigma-Aldrich). All cultures were maintained for 24 hours at 37°C with 5% CO₂.

After this incubation, plates were centrifuged at 800g for 5 minutes, and half of the supernatant volume was collected from all LDN and HDN conditions without disturbing the cell layer. This neutrophil-conditioned medium was then used in PBMC cultures. Thus, PBMC were cultured in RPMI without stimulus, in RPMI with stimulus, in neutrophil-conditioned medium (from either LDN or HDN) with stimulus, and in neutrophil-conditioned medium with stimulus and treated with nor-NOHA. Stimulation was performed using PMA (35 ng/mL) and Ionomycin (1 μg/mL) (Merk Millipore).

These cultures were incubated for an additional 24 hours at 37°C 5% CO₂, with Brefeldin A (BioLegend) added to all conditions 4 hours before incubation ended to block cytokine secretion. Following incubation, cells were stained with anti-CD3, anti-CD8, anti-CD4, anti-CD69, and anti-CD25 mAbs, as previously described. Samples were analyzed by flow cytometry using a BD FACS Canto II with FACSDiva software, and data were processed using FlowJo software.

### Statistical analysis

Statistical analysis was conducted using GraphPad Prism v9.5.0, and statistical significance was considered for p-values lower than 0.05. Comparisons between groups were performed using Kruskal-Wallis, Chi-Squared, Mann-Whitney, one-way ANOVA, or Friedman’s tests, as appropriate. A receiver operating characteristic (ROC) curve analysis using survival data from metastatic BC patients was performed to define a threshold for LDN frequency, aiming to minimize false negatives. This threshold was then applied to the progression analyses, survival curves, and the swimmer plot. Survival curves were analyzed using the Gehan-Breslow-Wilcoxon test. Data are presented as mean ± standard deviation (SD) unless otherwise stated.

## Results

### Low-density neutrophil accumulation in the blood of metastatic breast cancer patients is associated with a worse prognosis

Our previous study demonstrated that systemic low-density neutrophils (LDN) are increased in the blood of breast cancer (BC) patients compared to healthy individuals [16]. These cells were also associated with advanced disease and poor response to neoadjuvant chemotherapy in earlier stages, potentially leading to future recurrence and metastasis [16].

To further validate the significance of LDN in BC progression, we analyzed, by flow cytometry, the frequency of these cells within the peripheral blood mononuclear cells (PBMC), in three larger groups of individuals: healthy donors, non-metastatic BC patients, and metastatic BC patients. As expected, LDN were almost absent in the blood of healthy individuals compared to non-metastatic (p=0.0180) and metastatic BC patients (p=0.0003) (Figure 1A). Moreover, among BC patients, LDN levels were significantly higher in metastatic patients than in non-metastatic patients (p=0.0289) (Figure 1A).

**Fig. 1.**
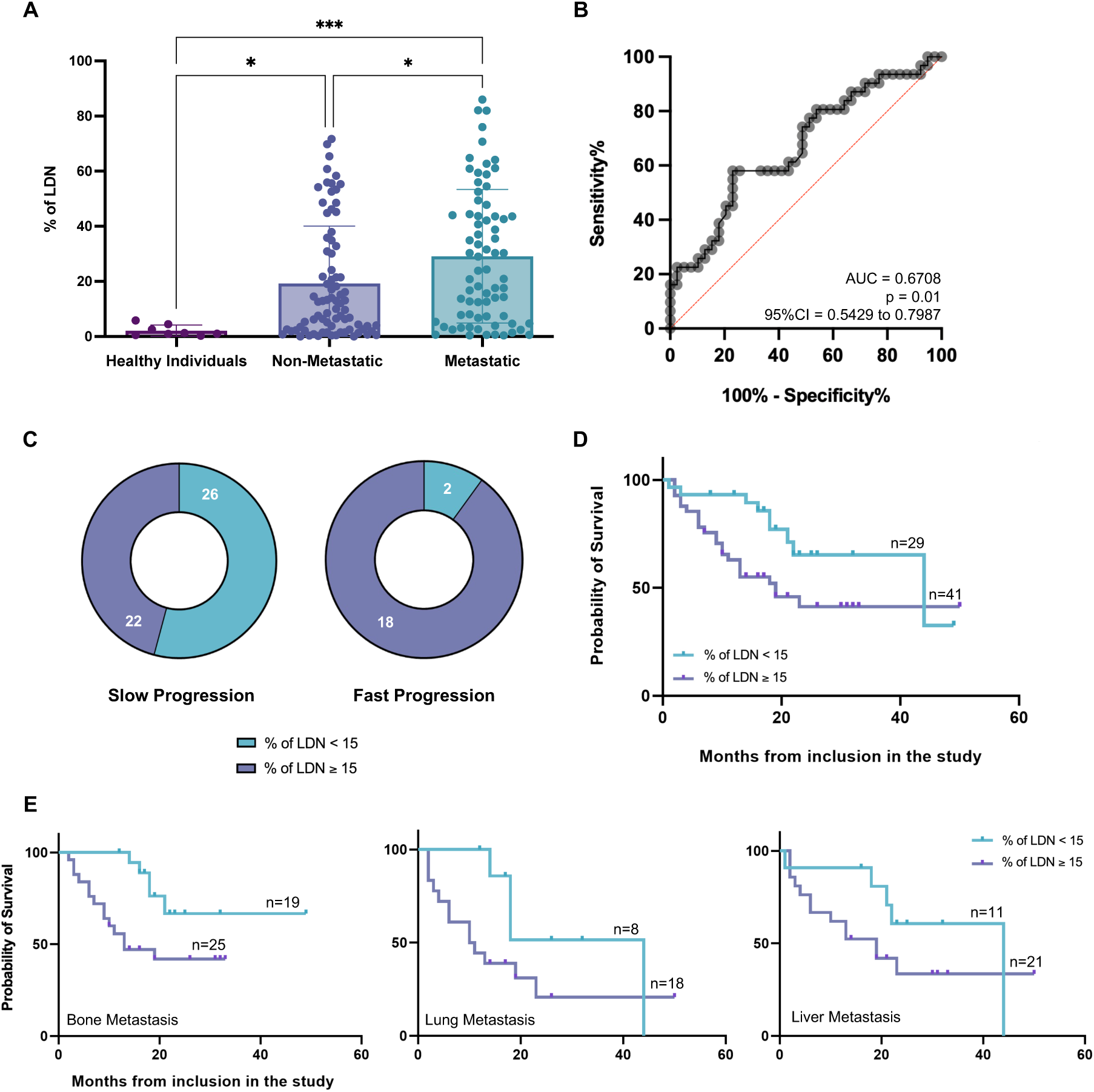
Low-density neutrophils accumulate in metastatic breast cancer and correlate with worse prognosis and reduced survival. A -. Percentage of low-density neutrophils (LDN) (assessed within the peripheral blood mononuclear cell (PBMC) layer) in the blood of healthy donors (plum bar, n=8), non-metastatic breast cancer (BC) patients (dark blue bar, n=79), and metastatic breast cancer (mBC) patients (teal bar, n=72). **B –** ROC curve evaluating the predictive performance of LDN frequency for patient prognosis (considering mBC patients’ survival data). The area under the curve (AUC) was 0.6708. The cut-off value obtained for the LDN percentage was 15% (70.97% sensitivity and 51.28% specificity, minimizing false negatives). **C –** Distribution of LDN levels in slow-progressing (n=48) and fast-progressing (n=20) mBC patients. Patients were categorized based on the percentage of LDN in their blood: LDN < 15% (blue) and LDN ≥ 15% (purple). **D –** Probability of survival of mBC patients exhibiting less than 15% (blue line) and 15% or more (purple line) of LDN in their blood. Hazard ratio: 0.4460 (95% CI, 0.22-0.90), *p=0.0104 (n=70). **E –** Probability of survival of mBC patients exhibiting less than 15% (blue line) and patients exhibiting 15% or more (purple line) of LDN in their blood, according to metastasis location: bone metastasis (Hazard ratio: 0.3087 (95% CI, 0.13-0.76), **p=0.0036, n=44); lung metastasis (Hazard ratio: 0.04342 (95% CI, 0.17-1.14), *p=0.0332, n=26); and liver metastasis (Hazard ratio: 0.5482 (95% CI, 0.21-1.40), p=0.0935, n=32). *p<0.05, ***p<0.001.

Notably, this increase in LDN frequency with disease severity occurs despite a reduction in the total frequency of neutrophils in the whole blood (Supplementary Figure S2), likely due to treatment-induced neutropenia in advanced disease. These findings highlight the involvement of LDN in BC progression and further suggest that this neutrophil subpopulation may exhibit greater resilience.

Additionally, we observed variability in LDN frequency across all BC subtypes, with no subtype showing a stronger correlation with LDN presence (Supplementary Figure S3).

Next, we investigated the prognostic value of LDN within metastatic BC (mBC) patients’ cohort. Using ROC curve analysis, we established a 15% LDN threshold that optimized sensitivity (70.97%) and specificity (51.28%) to minimize false negatives (Figure 1B). Notably, most mBC patients with rapid disease progression — defined by adverse clinical events or death within one year of their first blood collection for our study — had LDN levels above this threshold. In contrast, patients with slower disease progression exhibited LDN levels below 15% (p<0.001) (Figure 1C). Moreover, mBC patients with LDN levels above 15% had significantly shorter survival from the time of enrollment in our study compared to those with lower levels (<15%) (p=0.0104) (Figure 1D). Notably, this association was independent of these patients’ metastasis location. Patients with bone and lung metastases presenting higher levels of LDN (≥15%) in their blood at the time of their first blood collection for our study had reduced survival compared to those presenting lower LDN frequencies in their blood (p=0.0036 and p=0.0332, respectively), with a similar trend observed in patients with liver metastases (Figure 1E).

Altogether, these results reinforce the detrimental role of LDN in BC, independently of the subtype, suggesting their potential as prognostic biomarkers to identify metastatic patients at greater risk.

### Low-density neutrophils’ dynamics correlate with changes in the clinical status of metastatic breast cancer patients

To further clarify the clinical relevance of LDN and validate their value as biomarkers, we conducted a longitudinal study in our cohort of mBC patients, tracking LDN frequency over time in relation to disease status, whenever possible. The swimmer plot in Figure 2A illustrates individual follow-up periods, with each bar representing the duration of patient monitoring. Color segments denote LDN percentages at different time points, while symbols mark critical clinical events, including disease progression and treatment responses.

**Fig. 2.**
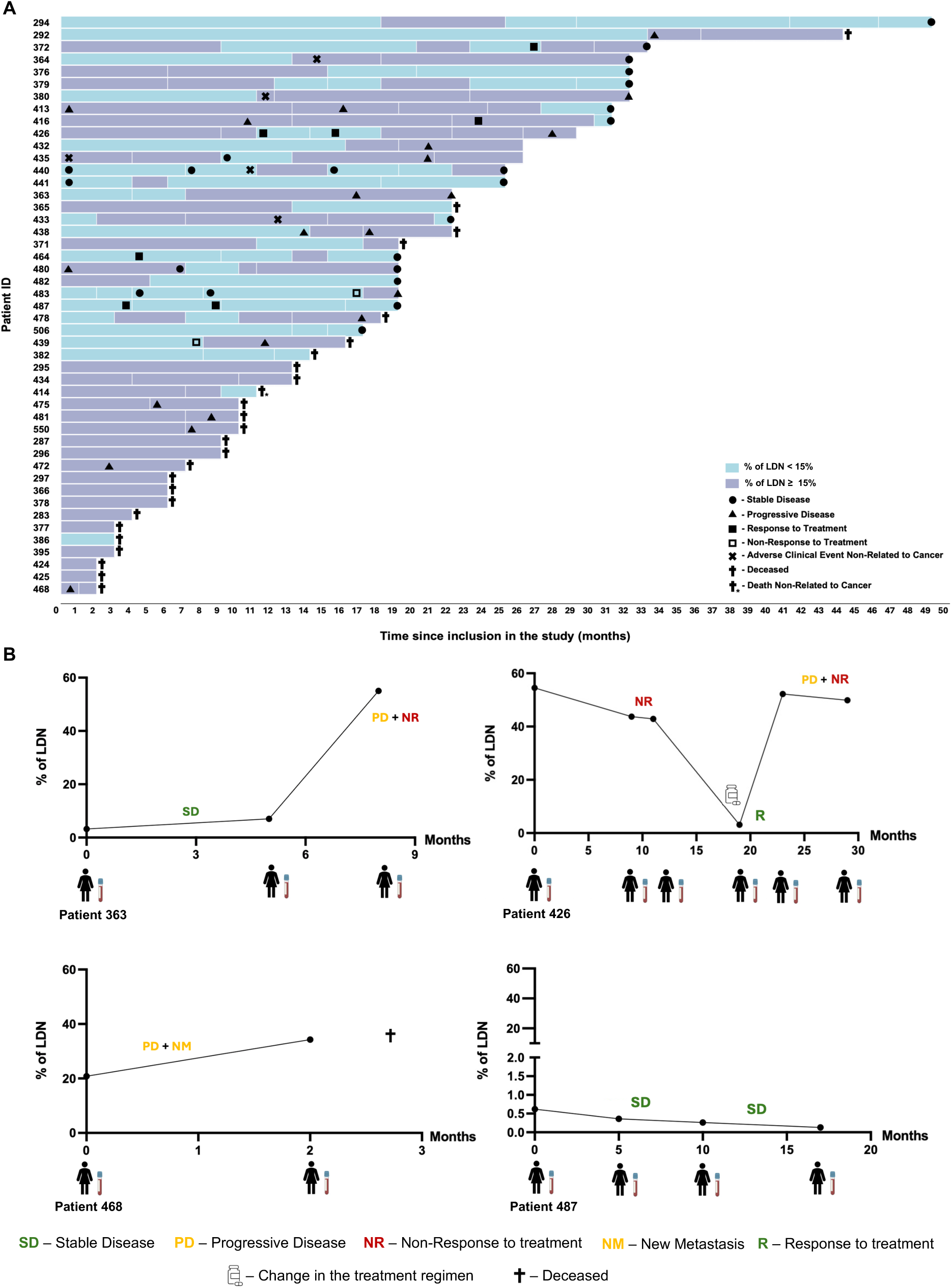
The dynamics of low-density neutrophils is associated with the clinical status of metastatic breast cancer. A –. Swimmer plot illustrating the percentage of low-density neutrophils (LDN) (assessed within the peripheral blood mononuclear cell (PBMC) layer) of 48 metastatic breast cancer (mBC) patients and respective relevant clinical events over time. Each bar represents a patient, with its length corresponding to the study enrolment period (in months). The colored segments reflect the LDN percentages in the blood, while symbols indicate relevant clinical events. **B –** Illustrative cases displaying longitudinal fluctuations in LDN percentages over time in relation to clinical outcomes for four specific mBC patients.

Patients with persistently high LDN levels (≥15%) frequently experienced rapid disease progression and adverse clinical events compared to those with lower LDN percentages. Conversely, individuals with fluctuating or consistently reduced LDN frequencies tended to exhibit prolonged disease stability and better treatment responses. While this association is not absolute, these findings suggest that LDN dynamics may reflect shifts in the clinical status of mBC patients and could serve as promising biomarkers for monitoring alterations in disease status, guiding clinical management.

To better illustrate these dynamics, we highlighted four specific cases (Figure 2B). Notably, sharp increases in LDN percentages were often followed by clinical events indicative of disease progression, including the development of new metastases, lack of treatment response, or shorter relapse intervals (Figure 2B; Patients 363, 426, and 468); whereas periods in which LDN remained low coincided with treatment response and prolonged disease stability (Figure 2B; Patients 363 and 487).

These findings underscore the potential of LDN as a real-time biomarker for tracking disease progression and predicting clinical outcomes in mBC. Monitoring LDN fluctuations could provide valuable insights into disease management, facilitating early detection of adverse events and aiding in therapeutic decision-making.

### Low-density neutrophils from metastatic breast cancer patients are highly immature cells and display enhanced pro-tumorigenic properties

Given the clinical relevance of LDN and their association with BC worse prognosis, we performed a phenotypic and functional characterization of LDN compared to high-density neutrophils (HDN), to further depict their distinct properties.

Flow cytometry analysis was performed to assess the percentage of LDN and HDN expressing key markers, including CD11b, CD66b, CD33, PD-L1, TLR4, MMP-9, MPO, VEGF, and elastase. CD11b and CD66b are adhesion molecules indicative of neutrophil activation [28], while CD33 is a marker of neutrophil immaturity, with its expression decreasing as cells mature [29]. PD-L1 is an immune checkpoint molecule that suppresses T-cell activity upon interaction with the PD-1 receptor [25]. TLR4 expression and activation are associated with prolonged neutrophil lifespan and correlate with the secretion of APRIL, a factor implicated in BC progression [30,31]. MMP-9, a matrix metalloproteinase, facilitates tumor invasion, metastasis, and angiogenesis by degrading the extracellular matrix [32]. Vascular endothelial growth factor (VEGF) is also a key promoter of neovascularization, supporting tumor growth [33]. Lastly, myeloperoxidase (MPO) and elastase are neutrophil-derived granular proteins involved in neutrophil extracellular traps (NETs) and indicators of neutrophil activation [34,35]. Additionally, MPO plays a crucial role in metastasis and immunosuppression [36,37].

Our analysis revealed a significantly increased percentage of LDN expressing CD33 (p<0.0001), PD-L1 (p<0.0001), TLR4 (p<0.0001), MMP-9 (p=0.0011), VEGF (p=0.0332) and MPO (p=0.0042) compared to HDN (Figure 3A). Collectively, these findings highlight that LDN from BC patients are immature neutrophils with distinct immunosuppressive and tumor-supporting characteristics.

**Fig. 3.**
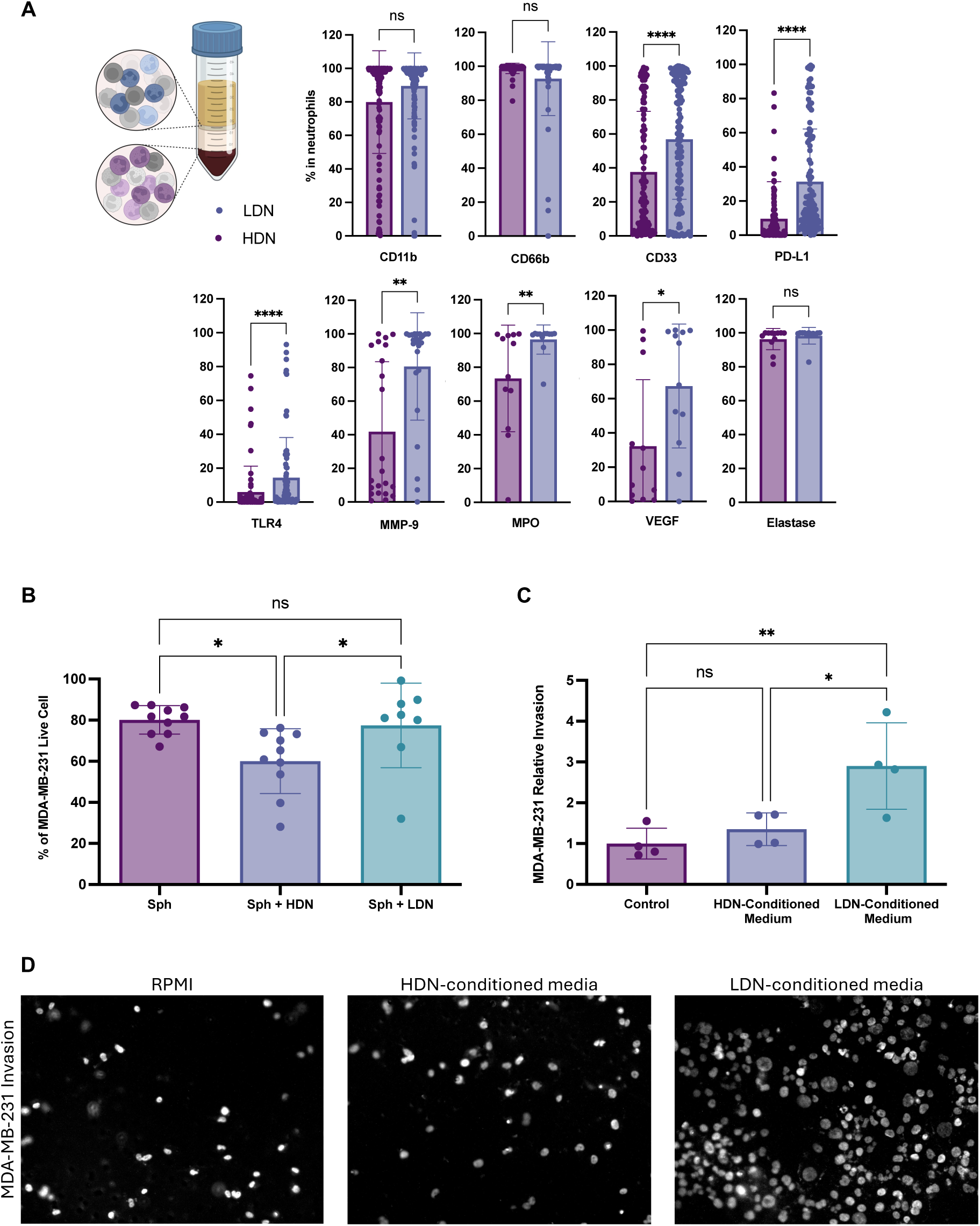
Low-density neutrophils are more immature cells with increased immunosuppressive and protumor features than high-density neutrophils. A -. Percentage of high-density neutrophils (HDN, plum bars) (assessed within the granulocyte layer) low-density neutrophils (LDN, dark blue bars) (assessed within the peripheral blood mononuclear cell (PBMC) layer) expressing CD11b (n=139), CD66b (n=69), CD33 (n=128), PD-L1 (n=116), TLR4 (n=75), MMP-9 (n=25), MPO (n=12), Elastase (n=12) and VEGF (n=12), assessed by flow cytometry. **B –** Percentage of live breast cancer (BC) cells in *in vitro* 3D cultures of MDA-MB-231 cell line alone (Sph, plum bar, n=10) or co-cultured with either HDN (Sph + HDN, dark blue bar, n=10) or LDN (Sph + LDN, teal bar, n=8). **C –** Relative invasion of MDA-MB-231 cells in Matrigel-coated transwells under different conditions: RPMI medium (control, plum bar, n=4), HDN-conditioned medium (dark blue bar, n=4), and LDN-conditioned medium (teal bar, n=4). **D –** Representative DAPI-stained images of MDA-MB-231 invasion through Matrigel-coated transwells under the same conditions as C. *p < 0.05, **p < 0.01, ***p < 0.001, ***p < 0.0001; ns = no significant difference.

To further explore the functional differences between these patient-derived neutrophil populations, we leveraged a 3D co-culture model previously established by our group [27]. We assessed the viability of a BC cell line following incubation with either LDN or HDN. While HDN significantly reduced BC cell viability (p=0.0109), consistent with their anti-tumorigenic phenotype, LDN failed to exert a cytotoxic effect on cancer cells (Figure 3B). The elevated expression of MMP-9, MPO, and VEGF in LDN corroborated their role in promoting tumor metastasis. This was confirmed through transwell invasion assays, in which BC cell lines were cultured with conditioned media from either LDN or HDN. BC cells cultured with LDN-conditioned medium exhibited a significantly enhanced ability to degrade and invade Matrigel-coated transwells, compared to those cultured with HDN-conditioned medium (p=0.0275) (Figures 3C and 3D). Notably, LDN-conditioned medium also led to increased BC cell proliferation, in contrast to medium from HDN cultures (p=0.0085) (Supplementary Figure S4). These results indicate that LDN from patients with advanced disease secrete soluble factors that play a critical role in supporting tumor cell proliferation and dissemination, underscoring their contribution to BC metastasis in alignment with the clinical observations.

Overall, these findings support the concept of LDN as key contributors to BC progression and highlight their potential as therapeutic targets to mitigate disease advancement.

### Low-density neutrophils impair ePector T lymphocyte function through soluble factors, including arginase

Our previous findings demonstrated that LDN can impair T lymphocyte activation when co-cultured [16]. Here, we aimed to clarify whether this effect is solely mediated by cell-cell interactions, for instance, through PD-L1/PD-1 – given the increased frequency of PD-L1-expressing LDN – or whether soluble factors released by LDN also contribute to T-cell dysfunction. To address this, we stimulated T lymphocytes in the presence of LDN- or HDN-conditioned media. T lymphocytes exposed to LDN-conditioned media exhibited significantly reduced activation, as indicated by a decrease in the expression of CD69 (p=0.0163) and CD25 (p=0.0066), two classical T-cell activation markers, and IFN-γ (p=0.0105), a key cytokine for T-cell mediated cytotoxicity (Figures 4A and 4B). These results suggest that LDN-released soluble factors are also mediators of immunosuppression.

**Fig. 4.**
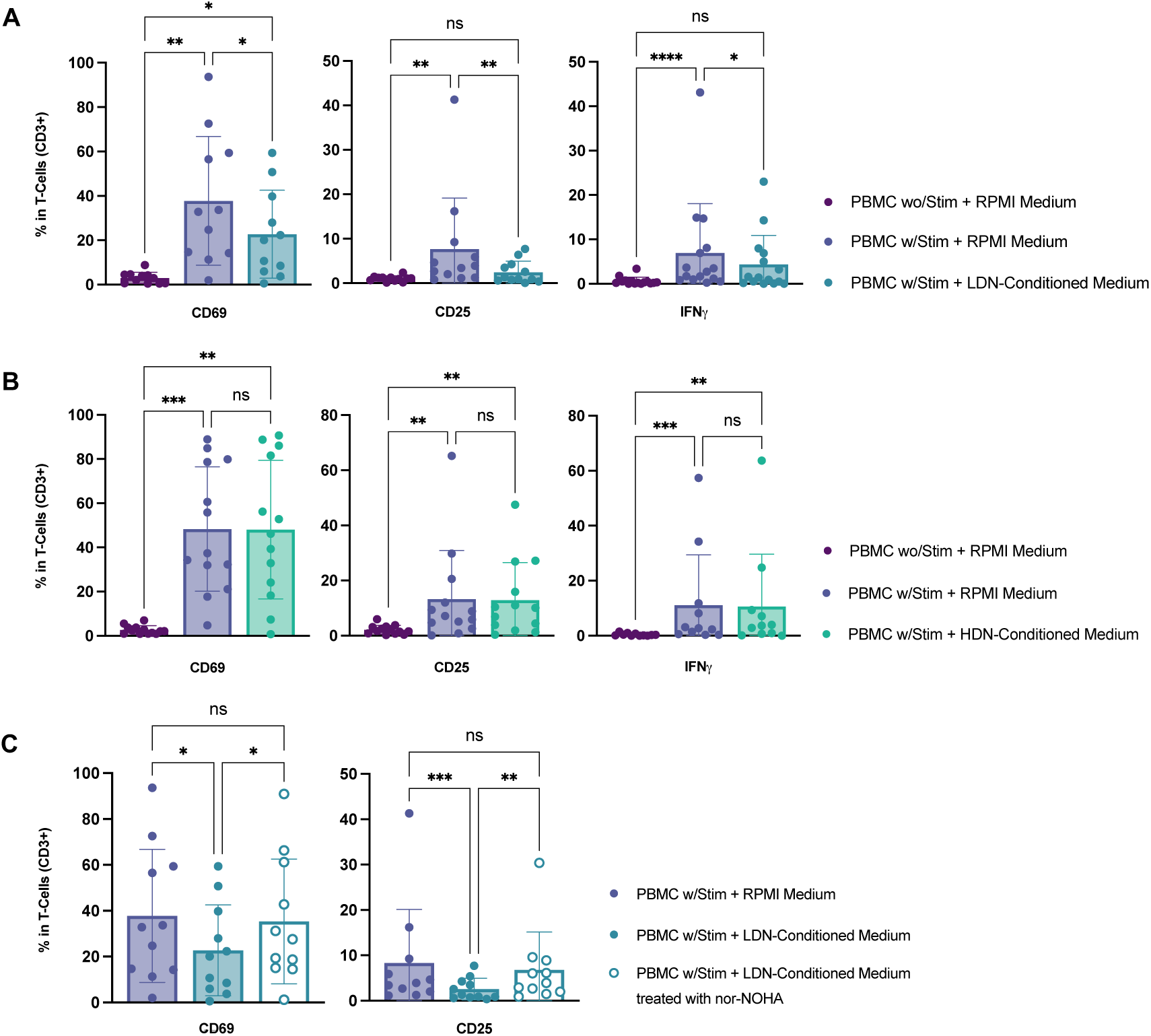
Low-density neutrophils contributed to ePector T lymphocytes’ dysfunction through soluble mediators, including arginase release. A –. Percentage of T-cells expressing the activation markers CD69 (n=11), CD25 (n=11), and IFNγ (n=11) within non-stimulated peripheral blood mononuclear cells (PBMC) (plum bars), stimulated PBMC, incubated in RPMI medium (dark blue bars) or stimulated PBMC exposed to LDN-conditioned medium (teal blue bars). **B –** Percentage of T-cells expressing the activation markers CD69 (n=11), CD25 (n=11), and IFNγ (n=11) within non-stimulated PBMC (plum bars), stimulated PBMC, incubated in RPMI medium (dark blue bars) or stimulated PBMCs exposed to high-density neutrophils (HDN)-conditioned medium (green bars). **C –** Percentage of T-cells expressing the activation markers CD69 (n=11) and CD25 (n=11) within stimulated PBMC (dark blue bars), exposed to LDN-conditioned medium (teal bars) or stimulated PBMC exposed to LDN-conditioned medium following LDN treatment with an arginase inhibitor (nor-NOHA) (blank bars). *p<0.05, **p<0.01, ***p<0.001, ****p<0.0001, ns – no significant difference.

A well-recognized soluble factor released by activated neutrophils and linked to T-cell dysfunction in several pathologies is arginase. This enzyme depletes arginine from the surrounding medium, thereby downregulating T-cell activation [38]. Notably, in the context of cancer, increased arginase expression is frequently associated with pro-tumorigenic N2 neutrophils [12,39].

To investigate the potential involvement of arginase in the immunosuppression specifically mediated by LDN-derived soluble factors in mBC patients, we performed the previously described T-cell stimulation assay in the presence of LDN-conditioned medium, this time adding nor-NOHA to the culture, a well-known arginase inhibitor [40]. Under these conditions, T lymphocytes achieved activation levels comparable to the control group, with no significant differences in CD69 and CD25 expression (Figure 4C). This indicates that the inhibitor was able to reverse, at least partially, the immunosuppressive effect of LDN-derived factors, suggesting a role for arginase in LDN-mediated T-cell dysfunction.

Altogether, these findings reinforce the immunosuppressive role of LDN in mBC, highlighting their ability to impair T-cell activation through the release of soluble factors.

## Discussion

Breast cancer (BC) is the leading cause of cancer-related mortality among women worldwide, accounting for approximately 700,000 deaths annually [1]. Metastasis poses the greatest challenge, with metastatic breast cancer (mBC) being responsible for the majority of these fatalities [2]. Indeed, despite the efforts to refine mBC management resulting and evidence of improvements in survival for patients with recurrent disease, 20-30% of early diagnosed BC patients still develop metastatic disease [2,41]. Currently, mBC patients mostly rely on conventional approaches aimed at prolonging life expectancy and improving their quality of life [3]. Nevertheless, resistance to the available treatment options remains a significant challenge, largely due to immunosuppression imposed by the tumor microenvironment (TME), which can influence cancer progression through intricate and context-dependent mechanisms [4,5]. Tumor-infiltrating immune cells, immunosuppressive soluble factors, and an altered extracellular matrix (ECM) collectively contribute to immunotolerance, suppress antitumor immunity, and facilitate BC progression and metastasis [42,43,44]. Among the immune components shaping the TME, neutrophils have gained attention for their dual role in cancer. While traditionally recognized as first-line defenders against pathogens, neutrophils, particularly low-density neutrophils (LDN), can adopt an immunosuppressive phenotype that supports tumor growth and metastasis [6,11,15,16,17]. In BC, these protumor neutrophils contribute to immune evasion by modulating T-cell responses, enhancing ECM remodeling, and promoting angiogenesis [16,32,45,46,47]. Understanding the mechanisms by which neutrophils, especially the circulating LDN, contribute to cancer progression and treatment resistance is crucial for improving patient outcomes.

This study provides compelling clinical evidence that highlights the significance of LDN, demonstrating through longitudinal follow-ups that fluctuations in LDN levels are closely associated with changes in clinical status. Our findings revealed that increased LDN percentages often preceded or coincided with adverse clinical events, including metastasis development and treatment resistance, suggesting their potential as real-time biomarkers for ongoing disease monitoring. Routine assessment of LDN levels could serve as an early indicator of disease progression, helping timely identification of high-risk patients, and aiding in clinical decision-making and treatment adjustments.

Moreover, our data reinforce the pro-tumorigenic role of LDN in human BC, demonstrating their ability to shape an immunosuppressive, invasive, and angiogenic TME. The elevated expression of matrix metalloproteinase-9 (MMP-9) and vascular endothelial growth factor (VEGF) in LDN highlights their role in facilitating ECM remodeling and metastasis promotion. MMP-9, a key matrix metalloproteinase, degrades structural barriers, enhancing cancer cell invasion and dissemination [32,48,49]. This aligns with our transwell invasion assay, where BC cells cultured with LDN-conditioned media exhibited significantly greater invasive capacity than those exposed to HDN-conditioned media. Additionally, the increased expression of VEGF in LDN suggests their involvement in neovascularization, a crucial process supporting tumor growth and metastatic spread [33,50]. These findings are consistent with previous studies linking MMP-9 and VEGF with poor prognosis in BC [51] as well as with observations in other cancers, such as gastric cancer [52,53], where these factors synergize to enhance tumor aggressiveness.

Furthermore, higher levels of myeloperoxidase (MPO) detected in LDN indicate another immunosuppressive mechanism and role in neutrophil extracellular trap (NET) formation, a process associated with metastasis and immune evasion [36,54]. This agrees with our previous observation, which showed increased MPO intensity in NETs produced by LDN compared to HDN [16].

Aligned with recent studies highlighting the role of Toll-like receptor 4 (TLR4) in BC progression [55,56], we observed that LDN exhibit higher TLR4 expression, than the normal neutrophils. TLR4 activation has been associated with prolonged neutrophil lifespan, and increased secretion of APRIL, a factor also implicated in BC development [30,31,57]. The observed increased expression of TLR4 on LDN compared to HDN may explain why LDN levels remain elevated despite treatment-induced neutropenia, which typically leads to a reduction in the overall neutrophil levels in patients’ blood. In accordance with this, we observed a decrease in total neutrophil counts in advanced disease, further emphasizing the endurance of LDN. Additionally, the TLR4-APRIL axis may amplify the pro-tumorigenic effects of LDN-derived MMP-9 and VEGF. It has been reported that heparan sulfate (HS) on the surface of BC cells activates TLR4 on neutrophils, triggering APRIL secretion [31]. In turn, neutrophil-derived APRIL binds back to BC cell HS, activating MAPK cascades (p38, ERK1/2, JNK1/2) and promoting proliferation [57]. Our findings, showing increased BC cell proliferation and invasion in response to LDN-conditioned media, align with this synergistic interaction between TLR4, MMP-9, and VEGF, potentially contributing to enhanced tumor growth and metastasis.

Collectively, these results suggest that LDN and their secreted factors not only facilitate tumor invasion but also establish a microenvironment favorable to tumor growth and immunosuppression. The interplay between TLR4, APRIL, and the key pro-tumorigenic factors (MMP-9, VEGF, MPO) highlights the potential of targeting LDN or their associated signaling pathways to simultaneously disrupt multiple tumor-promoting mechanisms, presenting a promising avenue for novel BC treatments. Additionally, the observed differential effects of neutrophil subsets on BC cell viability underscore the functional heterogeneity of neutrophil populations in the tumor microenvironment. While HDN significantly reduced BC cell viability, aligning with the proposed anti-tumorigenic N1 phenotype, LDN failed to exert cytotoxic effects, reflecting a pro-tumorigenic N2-like state [6,11,12,15,16,17]. This dichotomy highlights the complex interplay within the TME and may explain the challenges encountered in neutrophil-targeted cancer therapies. The cytotoxic effect of HDN could be attributed to mechanisms such as the release of TRAIL or reactive oxygen species, protease activity, or direct cell-cell interactions, whereas the lack of LDN cytotoxicity might stem from altered effector functions or immunosuppressive properties [58,59]. These findings suggest that therapeutic strategies aimed at enhancing HDN function or converting LDN to an HDN-like phenotype could be promising. Beyond their role in creating a pro-proliferative and invasive TME, our results confirm that LDN also exert immunosuppressive effects on T-cells. Previously, we demonstrated that LDN impair T-cell responses through direct cell-cell interactions [16]. Here, we further show that T-cells stimulated under the influence of LDN-conditioned media exhibit a significant reduction in their activation markers (CD69, CD25) and IFN-γ production, highlighting the pivotal role of LDN-derived soluble factors in mediating T-cell dysfunction.

Our investigation into the mechanisms underlying LDN-mediated immunosuppression identifies arginase as a key contributor. The ability of the arginase inhibitor nor-NOHA to restore T-cell function strongly implicates this enzyme in LDN-driven immunosuppression and underscores its relevance as a potential therapeutic target. This finding aligns with research linking increased arginase activity to pro-tumorigenic N2 neutrophils in cancer [37,38]. While arginase inhibition is not a novel concept in cancer treatment [40,60,61], its clinical success has been limited [62]. Nonetheless, our results, emphasize the need to reconsider its therapeutic relevance within the appropriate clinical context. Targeting arginase is likely to be more effective in individuals with a high frequency of cells overexpressing this enzyme, particularly those with elevated LDN levels. However, most clinical trials evaluating arginase inhibitors have been conducted without stratifying patients based on relevant biomarkers, which may have contributed to their limited efficacy. Therefore, refining the design of clinical trials is essential, incorporating LDN as a biomarker to better identify the patients who could truly benefit from arginase-targeted therapies.

## Conclusions

Our findings highlight the critical role of LDN in BC aggressiveness (Figure 5). To our knowledge, we provide the first longitudinal evidence demonstrating the potential of LDN for monitoring BC progression. Their pro-tumorigenic and immunosuppressive properties, coupled with their increased prevalence in metastatic disease, underscore their value as both biomarkers and therapeutic targets. However, it is important to recognize that LDN comprise heterogeneous subsets, not all of them contributing equally to cancer progression. Indiscriminate targeting could disrupt essential immune functions, emphasizing the need for more refined studies.

**Fig. 5.**
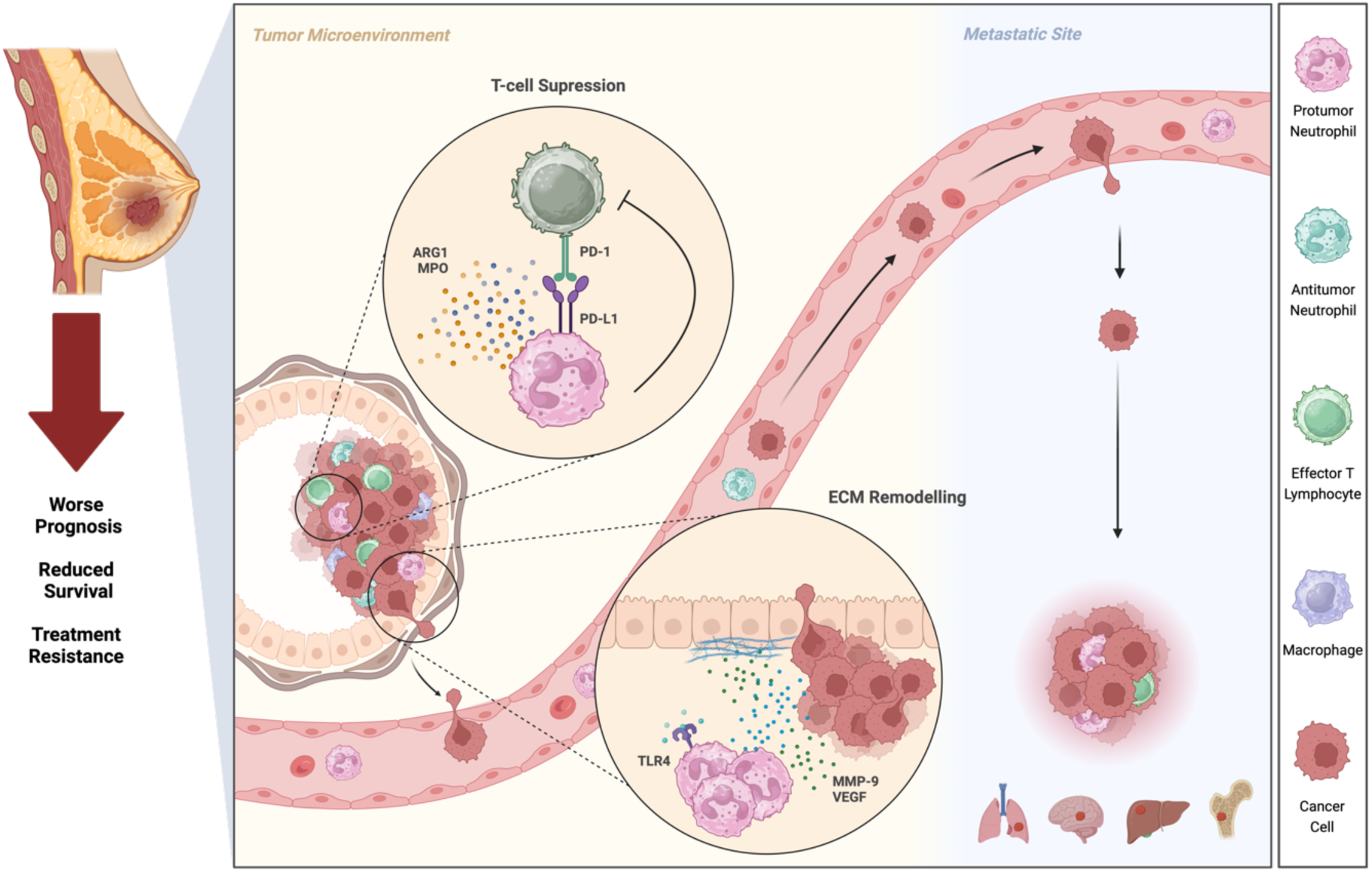
Low-density neutrophils promote breast cancer progression through immunosuppression and protumor activity. Unlike the antitumor high-density neutrophils (HDN), low-density neutrophils (LDN) are protumor cells that contribute to breast cancer (BC) progression by suppressing immune responses and enhancing tumor aggressiveness. Although LDN circulate systemically, they can be recruited to the tumor microenvironment, where they may suppress T-cell function both directly — via cell–cell interactions mediated by PD-L1, which is highly expressed in LDN — and indirectly, through soluble factors such as arginase (ARG1) and myeloperoxidase (MPO). Additionally, LDN promote tumor proliferation, invasion, and metastasis by secreting matrix metalloproteinase-9 (MMP-9) and vascular endothelial growth factor (VEGF). Toll-like receptor 4 (TLR4) expression is also markedly elevated in LDN, potentially contributing to tumor progression by inducing APRIL secretion. The accumulation of LDN with these characteristics in the blood of BC patients may underlie their association with treatment resistance, poor prognosis, and reduced survival.

Future research should focus on further characterizing neutrophil heterogeneity in BC, identifying specific biomarkers that distinguish pathological subsets from homeostatic neutrophils. This will facilitate the development of precision therapies that selectively modulate protumor neutrophils while preserving beneficial immune functions. The identification of these biomarkers will not only enhance our ability to target specific neutrophil subsets but also improve patient stratification for more personalized treatments. This approach holds great promise for boosting the efficacy of BC therapies and ultimately improving patient outcomes.

## Abbreviations

BC: Breast Cancer
ECM: Extracellular matrix
HDN: High-density neutrophils
LDN: Low-density neutrophils
mBC: Metastatic breast cancer
NETs: Neutrophil extracellular traps
PBMC: Peripheral blood mononuclear cells
TME: Tumor microenvironment

## Acknowledgments

We thank all breast cancer patients who generously agreed to participate in this study. We are also grateful to the dedicated nurses and oncologists from the participant hospitals, whose collaboration was fundamental for the collection of patient blood samples and clinical data. We would also like to thank the Flow Cytometry and Cell Culture Facilities of NOVA Medical School. Our acknowledgemnets are also extended to the support given by the Research Unit iNOVA4Health and by the Associated Laboratory LS4FUTURE. Some figures in this work were created using BioRender.com.

## Declarations

### Authors’ contributions

BFC and DG contributed equally to this work. Both conducted all the experiments, analyzed, and interpreted the data, performed the statistical analysis, assembled all the figures, and wrote the manuscript. RS and IB helped in the execution of flow cytometry and cell culture experiments. SB, TM, MV, CXS and SCF were responsible for selecting the patients, collecting the samples and clinical data, and contributing also to the scientific discussion. AJ contributed to scientific discussion. MGC designed and supervised the study, contributed to data interpretation, and participated in writing the manuscript. All the authors revised and approved the final manuscript.

### Ethics approval and consent to participate

Written informed consent was obtained from all individuals for the collection of biological samples and clinical data. The study complied with the ethical principles outlined in the Declaration of Helsinki and received approval from the Ethical Committees of HFF, ULSAR, HVFX, HSM, HCD, IPO, and NOVA Medical School.

### Consent for publication

Not applicable

### Availability of data and materials

No datasets were generated or analyzed during this study.

