## Supplementary Material for "Neutrophils matter: New clinical insights on their role in the progression of metastatic breast cancer"

### Table of Contents

Supplementary Figure S1 – Flowchart of the breast cancer patients enrolled in this study. 1

Supplementary Figure S2 – Low-density neutrophils' frequency increases despite of overall decrease in total neutrophils with breast cancer progression. 1

Supplementary Figure S3 – The frequency of low-density neutrophils in breast cancer patients shows similar variability across all subtypes. 2

Supplementary Figure S4 – Low-density neutrophils-conditioned medium enhanced breast cancer proliferation. 2

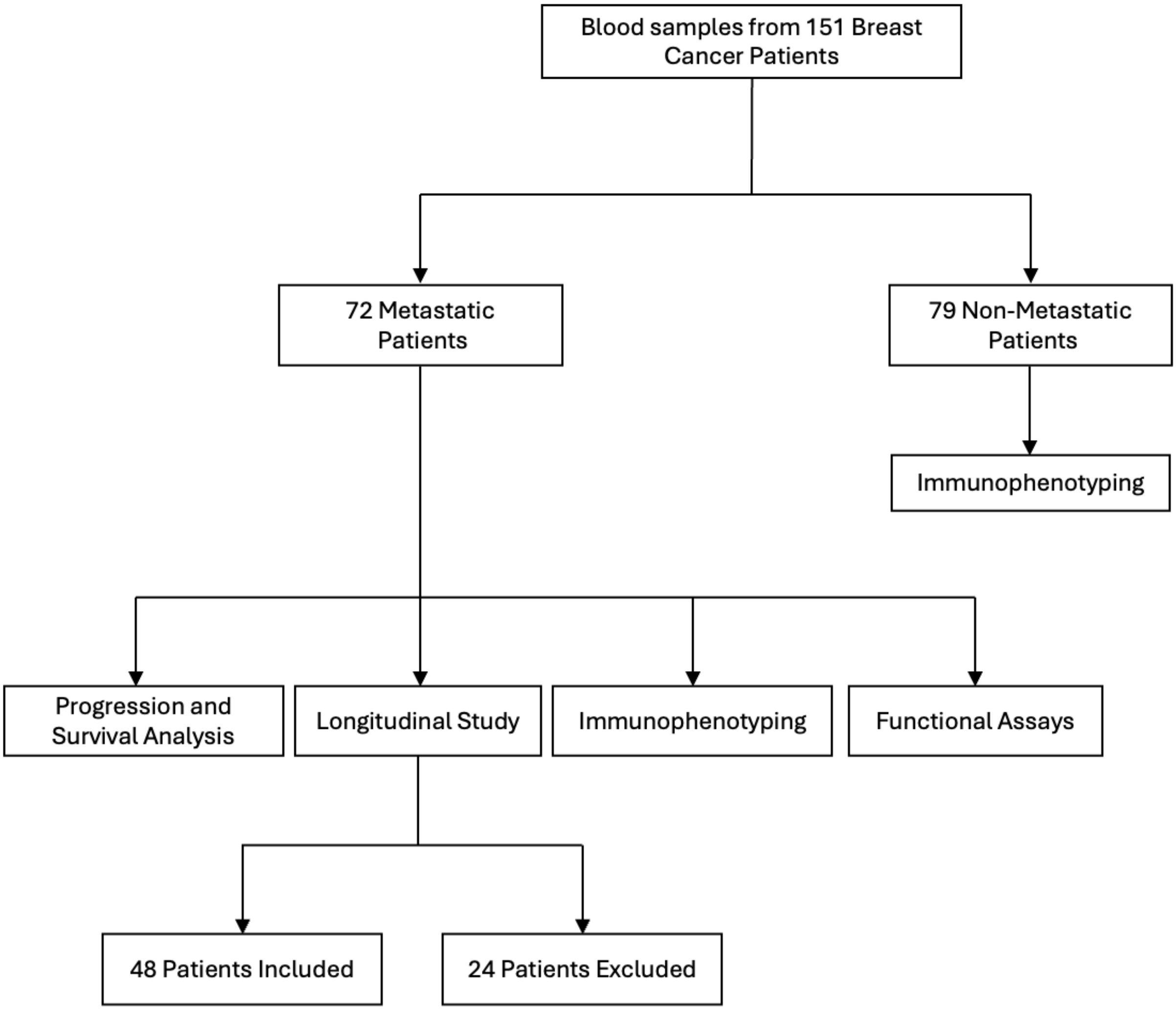


**Supplementary Figure S1 – Flowchart of the breast cancer patients enrolled in this study.** Blood samples from 151 breast cancer patients were analyzed, including 72 metastatic and 79 non-metastatic cases. The frequency of low-density neutrophils and high-density neutrophils as well as their respective immunophenotype were assessed in both cohorts. The neutrophils derived from metastatic patients were further used for progression and survival analysis, to conduct a longitudinal study, and for functional assays. For the longitudinal study, 48 of the 72 metastatic patients were included. The remaining 24 patients were excluded due to insufficient clinical data, widely spaced follow-up samples, or hospital transfer.

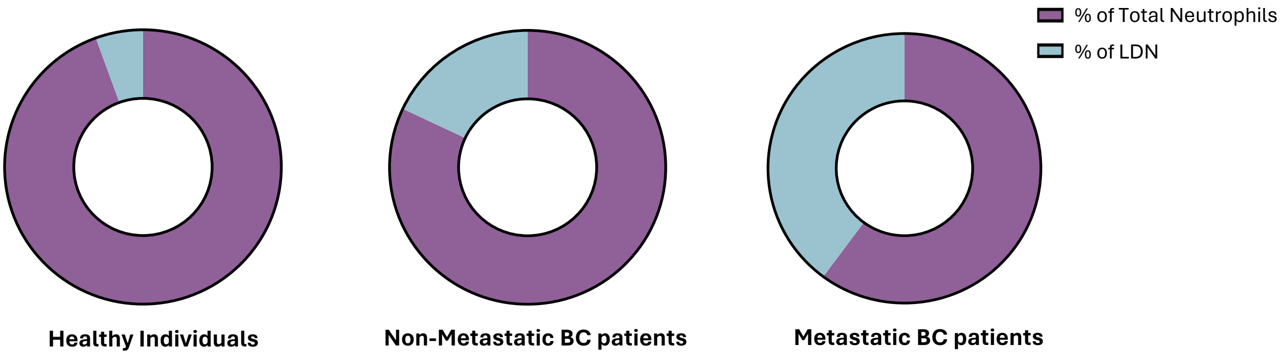


**Supplementary Figure S2 – Low-density neutrophils' frequency increases despite of overall decrease in total neutrophils with breast cancer progression.** Illustration of the distribution of total neutrophils in the whole blood and low-density neutrophils (LDN) within the peripheral blood mononuclear cell layer of healthy individuals (n=8), non-metastatic (n=79) and metastatic breast cancer (BC) patients (n=72).

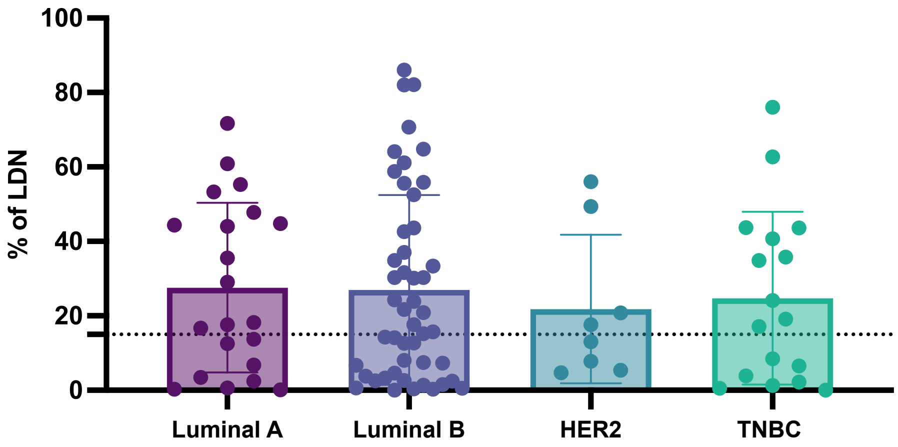


**Supplementary Figure S3 – The frequency of low-density neutrophils in breast cancer patients shows similar variability across all subtypes.** Percentage of low-density neutrophils (LDN) in the blood of breast cancer patients, categorized by subtype. The dotted line represents the reference threshold for elevated LDN levels (≥15%), as established using ROC curve analysis represented in Figure 1B. Luminal A (plum bar, n=21), Luminal B (dark blue bar, n=48), HER2 (teal bar, n=8), Triple-Negative (TNBC, green bar, n=17).

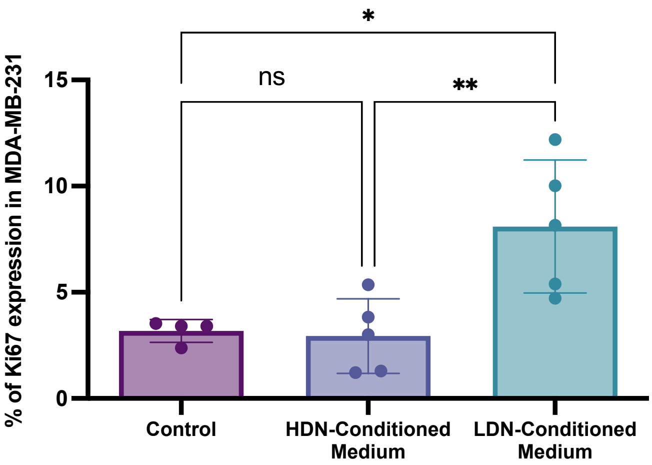


**Supplementary Figure S4 – Low-density neutrophils-conditioned medium enhanced breast cancer proliferation.** Percentage of Ki67-expressing MDA-MB-231 cells cultured in RPMI medium (plum bar, n=4), exposed to high-density neutrophils (HDN)-conditioned medium (dark blue bar, n=5) or low-density neutrophils (LDN)- conditioned medium (teal bar, n=5). *p<0.05, **p<0.01, ns – no significant difference.
